# NIMAA: an R/CRAN package to accomplish NomInal data Mining AnAlysis

**DOI:** 10.1101/2022.01.13.475835

**Authors:** Mohieddin Jafari, Cheng Chen, Mehdi Mirzaie, Jing Tang

## Abstract

**Summary:** Nominal data is data that has been “labeled” and can be designated into a number of non-overlapping unordered groups. The analysis of this type of data is often trivial because it is not feasible to conduct extensive numerical methods on this type of data. Graphs or networks, on the other hand, are comprised of sets of nodes and edges that can also be considered as nominal variables. By integrating graph theory and data mining approaches, we offer the R package NIMAA to define a nominal data-mining pipeline to explore more information. Using nominal variables in a dataset, NIMAA provides functions for constructing weighted and unweighted bipartite graphs, analysing the similarity of labels in nominal variables, clustering labels or categories to super-labels, validating clustering results, predicting bipartite edges by missing weight imputation, and providing a variety of visualization tools. Here, we also indicated the application of nominal data mining in a biological dataset with well-riched nominal variables.

**Availability:** NIMAA’s official release and the beta update are available on CRAN and Github, respectively. URLs: https://CRAN.R-project.org/package=NIMAA and https://github.com/jafarilab/NIMAA

**Contributions:** MJ conceived the study and developed the models, MJ and CC adopted and implemented the methods, MM improved the methods, JT provided the funding, MJ, CC, MM and JT wrote the paper.

## 1 Introduction

By expanding the concept of precision medicine, we progressively encounter to larger datasets with more features as well as nominal variables such as patients, cell lines, treatments and bio-annotations (Bhinder *et al.*, 2021; Josephson and Wiebe, 2021; Jafari *et al.*, 2021b). While the majority of data mining and machine learning techniques focus on numerical and ordinal variables, nominal data are frequently overlooked when computing similarities and characterizing labels. Numerous numerical analyses are invalid in this circumstance, as the mode is the only legitimate measure of central tendency for nominal data, and the appropriate statistical analysis is frequency testing, e.g., Chi-square tests to examine the proportion or frequency of labels in contingency table (Wu, 2017; Suich and Turek, 2003).

Consider a dataset with various chemical molecules tested on several patient-derived samples and a dataset containing multiple natural products with the molecular ingredient details. If exploring the pairwise similarity of patients or natural products are desired, there is an approach, for instance, to utilize correlation analysis on gene expression or metabolite concentration, respectively. However, in case that there are only the binary labels of potent chemicals on patients or multi-labels of ingredient availability in natural products, it is unclear whether further information might well be mined to explore similarities. On the other hand, precision medicine has advanced in recent years due to the development of numerous network-based tools and graph models (Jafari *et al.*, 2020; Turanli *et al.*, 2018). Graphs are composed of two distinct sets of nodes and edges, each of which includes labeled values. Additional sets of entities, such as the weight set in a weighted graph or several node sets in multipartite graphs, assist us in providing a more stringent graph model for a specific phenomenon. Leveraging these notions, we previously proposed using multipartite graphs to investigate pairwise similarities in nominal variables, with a particular emphasis on drug discovery research (Jafari *et al.*, 2020, 2021a). Here, we have proposed NIMAA, an open-source R package, that contains a set of functions that can be used to implement a series of bipartite graph-based data mining methods. Numerous tasks, ranging from the original nominal data input to the validation of the clustering findings employing prior knowledge, as well as imputation/prediction of missing data, can be accomplished. Similarly, validation and ranking of the imputation results are also included. Additionally, numerous visualization steps are addressed to support users in perceiving data structures. The overview flowchart of NIMAA is shown in Figure 1A.

**Fig. 1.**
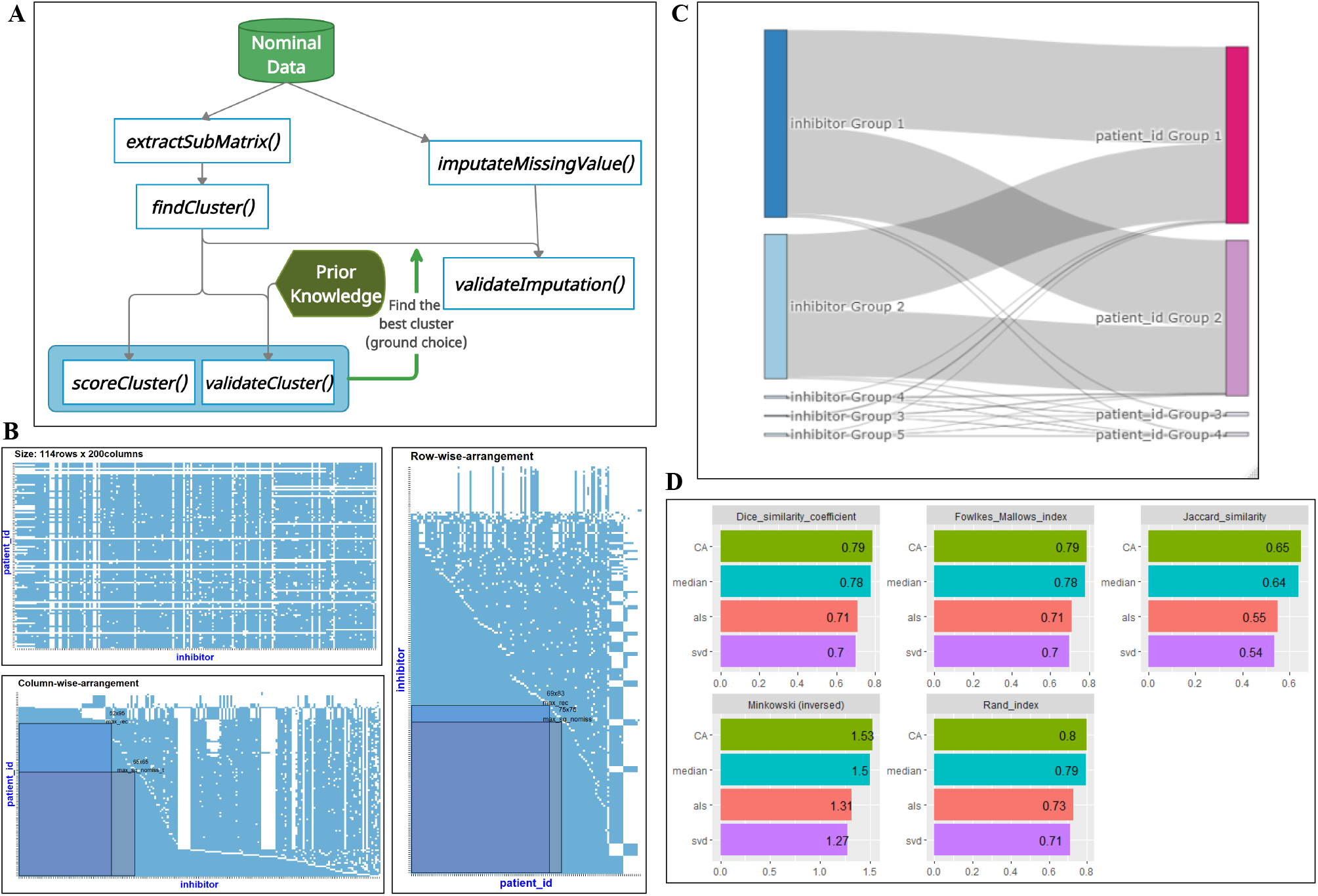
(A) Overview flowchart of NIMAA pipeline providing major functions, (B) An example dataset with a mixture of non-missing and missing values (blue and white area, respectively). The original data matrix, column-wise arrangement, and row-wise arrangement are shown with the highlighted large submatices of non-missing values with different sizes (C) The sankey plot of the bipartite graph to show the relationship of clusters found in the projected networks, and (D) The validation result of some imputation methodss. CA: Correspondence Analysis, als: alternating subspace SVD algorithm, svd: singular value decomposition

## 2 Implementation

### 2.1 Bird’s eye survey of data

A dataset with at least two nominal variables is taken as input data to reconstruct a bipartite network. This network can also be weighted by any optional numeric variables corresponding to pairs of nominal values. Therefore, the input data is a table consisting of two columns of nominal data plus one column of numerical data (optional), which is subsequently transformed into an incidence matrix (bi-adjacency matrix) by the `plotInput()` function. The heatmap plot of the matrix also provided to have a bird’s eye view of the non-missing values topology of the dataset, i.e., modularity and nestedness of non-missing values. The original input data can also be transformed into an `igraph` graph object by the function `plotBipartite()`, which also can plot the bipartite graph of original data.

### 2.2 Non-missing submatrix extraction

In the case of a weighted bipartite graph, downstream analysis should be performed on the dataset with the fewest missing values to avoid sensitivity issues of clustering-based methods. Therefore we use the `extractSubMatrix()` function to rearrange the original matrix to find the largest possible submatrices without missing values. Finding the best (largest) matrix without missing values has been shown to be an NP-hard problem (Bartholdi III, 1982), and this function will obtain submatrices in several ways as close to the largest submatrix (Figure 1B), see the package vignette for details of this function.

### 2.3 Analysis of projected networks

By transforming an incidence matrix into a bipartite graphs, clustering analysis of the projected networks describes the group of similar vertices in each vertex set based on similarity of neighbours in the bipartite graph. In a weighted bipartite graph, the pairwise similarity is enhanced by the weight magnitude. The whole process is done using the `findCluster()` function, which is further supplied with clustering quality analysis by returning *Silhouette*, *Coverage*, and *Modularity* metrics. If the dataset was supplemented with prior knowledge of categorization of labels, then it is possible for the user to use it as additional input. In this case, after completing the clustering analysis, two extra validation metrics, i.e., *Jaccard similarity coefficient* and *Rand index* is also available.

### 2.4 Bipartite edge prediction by weight imputation

The function `imputeMissingValue()` is able to impute missing weight values in a variety of ways, which can also be considered as a prediction of missing edges in a bipartite graph. It covers the classic numerical imputation functions such as mean, median, CA, and als. Subsequently, it will cluster the projected network of imputed incidence matrix exactly according to the input clustering parameters obtained from submatrix with non-missing values. To select the best imputation method, NIMAA provides comparison of the similarity analysis of non-missing submatrix with imputed matrix by returning *Sørensen-Dice coefficient*, *Fowlkes-Mallows index*, *Jaccard similarity*, *Minkowski index*, and *Rand index*.

## 3 Case study

Four exemplary datasets are available in NIMAA to cover all types of inputs including weighted, unweighted, modular and nested bipartite graphs. In a case study on the *beatAML* data (Tyner *et al.*, 2018), two nominal variables, namely inhibitors and patient identifiers, are selected, where the median of the corresponding drug responses are determined (i.e., edge weight). Firstly, the original incidence matrix is shown to perceive the amount and topology of non-missing values (Figure 1B). Secondly, to select the largest submatrix with non-missing values, column-wise and row-wise extraction were investigated. Notably, user selection may be influenced by biological reasoning (e.g., focusing on patients or on inhibitors) in addition to the size of the submatrix. Then, we perform a clustering analysis on the projected network of inhibitors and patient ids and visualize it in a bipartite graph to uncover cluster relationships (Figure 1C). Meanwhile, identification of the best clustering method can be done based on the cluster quality assessment. Finally, the best clustering method are used to predict bipartite edges and to evaluate the imputation results (Figure 1D).

## 4 Conclusion

NIMAA package provides a comprehensive set of pipeline to perform nominal data mining using graph theory. It can effectively uncover label relationships of each nominal variable according to pairwise association with other nominal variables. Edge prediction based on weight imputation is another application of NIMAA to explore local and global similarities within nominal variables. NIMAA is able to score and compare the results between pre- and post-imputation by utilizing internal and external (prior knowledge) assessment measures. This pipeline is strengthened by visualization and validation criteria that determine the optimal clustering method. This pipeline can be applied to a variety of nominal variables, regardless of the numerical values to quantify the relationship. Additionally, it was recently demonstrated that the NIMAA pipeline supports precision medicine particularly in network pharmacology by identifying synergistic medication combinations, patient resistance exploration, and clinical drug effects (Jafari *et al.*, 2021a). Taken together, the NIMAA pipeline is expected to facilitate the nominal data mining analysis not only in precision medicine, but also in more general applications like marketing and social science.

